# Immune Transcriptional Signatures Across Human Cardiomyopathy Subtypes: A Multi-Cohort Integrative Computational Analysis

**DOI:** 10.64898/2026.03.10.710912

**Authors:** Babatunde Adegboyega, Benson Okorie, Phillip Courage

## Abstract

**Background:** Heart failure, arrhythmia, and sudden cardiac death are common outcomes of cardiomyopathies, which are molecularly diverse heart muscle disorders marked by structural and functional myocardial dysfunction. The lack of sensitive molecular biomarkers that precede overt physiological deterioration makes early diagnosis difficult despite advancements in imaging and clinical classification. The immune transcriptional landscape across cardiomyopathy subtypes is still poorly understood, despite growing evidence linking both innate and adaptive immune dysregulation, such as macrophage activation and T-cell and inflammatory cytokine networks, as active contributors to myocardial remodelling and disease progression.

**Methods:** We performed a multi-cohort integrative transcriptomic analysis of 1,068 cardiac tissue samples from five publicly available GEO datasets (GSE57338, GSE5406, GSE36961, GSE141910, GSE47495) spanning dilated, ischemic, hypertrophic, and peripartum cardiomyopathy. Using a fully scripted R and Python pipeline, we conducted differential expression analysis (limma), immune cell deconvolution (xCell), pathway enrichment (clusterProfiler), weighted gene co-expression network analysis (WGCNA), and regularised machine learning classification (LASSO, Random Forest). Cross-dataset validation was performed in two independent cohorts on different microarray platforms.

**Results:** Differential expression analysis identified 43 primary DEGs (FDR < 0.05, |log2FC| > 1.0), revealing a coherent immune-fibrotic program characterized by loss of anti-inflammatory macrophage markers (CD163, VSIG4), complement dysregulation (FCN3), innate interferon activation (IFI44L, IFIT2), and ECM remodelling (ASPN, SFRP4, LUM). xCell deconvolution identified coordinated depletion of adaptive immune populations in failing myocardium. WGCNA defined a fibrosis hub module (brown; CTSK, SULF1, SFRP4) and an immune collapse module (turquoise; MYD88, TNFRSF1A, LAPTM5). A nine-gene LASSO classifier achieved a cross-validated AUC of 0.986, with HMOX2 as the top-discriminating feature, implicating ferroptosis in cardiomyocyte death. Cross-platform validation in an independent HCM cohort (GSE36961) demonstrated a directional concordance of 34.9%.

**Conclusions:** This study defines a reproducible immune-fibrotic transcriptional signature of human cardiomyopathy, nominates HMOX2 and ferroptosis as central pathomechanisms, and provides a validated nine-gene biomarker panel for future translational investigation.

## 1. Introduction

Cardiomyopathies are groups of heart muscle disorders that show structural and functional problems. These issues often lead to heart failure, arrhythmias, and sudden cardiac death (Ciarambino *et al.,* 2021; Precone *et al.,* 2019). Even with improvements in imaging, clinical classification, and molecular testing, detecting cardiomyopathy early is still a big challenge. This is particularly true for people without symptoms or those showing subtle signs of changes in the heart muscle. One key reason for the delayed diagnosis is the lack of early genetic markers that can predict disease onset before noticeable physiological problems arise (Ciarambino *et al.,* 2021; Brieler *et al.,* 2017). Current diagnostic methods mostly depend on observable traits like ventricular dilation, wall thickening, or reduced ejection fraction—indicators that typically show up only after significant damage has happened. There is a pressing need for molecular markers that reflect the earliest signs of stress on heart cells, immune responses, and changes in the heart structure (Ciarambino *et al.,* 2021; Bozkurt *et al.,* 2016). Recent studies suggests that cardiomyopathy is not just a mechanical or metabolic issue; it is increasingly seen as a complex disease involving molecular and immune factors. Research has indicated that both innate and adaptive immune pathways, particularly macrophage activation, T-cell signaling, and inflammatory cytokine networks, are crucial in heart muscle changes, scarring, and blood vessel issues. Genetic variations that influence immune signaling, such as mutations in TNNT2, LMNA, MYH7, and inflammatory regulators, have been linked to susceptibility to cardiomyopathy and varying clinical outcomes (Rao *et al.,* 2025; He *et al.,* 2021). However, most studies have focused on immune markers seen in advanced stages, leaving the early immune-related gene activity largely overlooked. Gaining insights into these initial changes in gene expression and genetic variations could help identify early biomarkers that are valuable for diagnosis and predicting health outcomes.

Computational genomics has become essential in contemporary cardiovascular research. High-throughput transcriptomic platforms, including microarrays and RNA-seq, have produced extensive gene expression datasets that capture disease-associated molecular alterations across diverse patient populations. Integrative multi-cohort approaches that incorporate differential expression analysis, immune deconvolution, gene co-expression network analysis, and machine learning classification demonstrate significant potential for identifying robust biomarker signatures (Mukherjee *et al.,* 2025; Zheng *et al.,* 2025; Casamassimi *et al.,* 2017). However, most existing cardiomyopathy transcriptomics studies are limited to single datasets and specific disease subtypes, which restricts the generalizability of their conclusions. The availability of multiple publicly archived datasets encompassing dilated, hypertrophic, and peripartum cardiomyopathy now enables systematic integrative analyses that were previously unattainable (Taylor *et al.,* 2023).

Despite improvements in cardiac transcriptomics, several important gaps still exist. First, most published studies look at only one cardiomyopathy subtype, which reduces understanding of shared and specific immune mechanisms. Second, immune cell composition has seldom been systematically analyzed from bulk transcriptomic data in cardiomyopathy, leaving the cellular basis of transcriptional changes poorly defined. Third, reproducible machine learning classifiers based on immune gene signatures have not been created for distinguishing cardiomyopathy subtypes and validated with independent cohorts. Fourth, the consistency of differential expression results across microarray platforms and disease subtypes has not been widely assessed. These gaps hinder the ability to translate transcriptomic findings into clinically useful biomarkers.

To address these issues, this study integrates transcriptomic data from 1,068 cardiac tissue samples across four publicly available GEO datasets that cover dilated, ischemic, hypertrophic, and peripartum cardiomyopathy. Using a multi-step analytical pipeline in R and Python, we conduct differential expression analysis, immune cell deconvolution, pathway enrichment, weighted gene co-expression network analysis, and regularized machine learning classification. Cross-dataset validation is used to evaluate the consistency of immune signatures across cardiomyopathy subtypes and independent groups. This study seeks to describe the shared and specific immune transcriptional landscape of human cardiomyopathy, identify a reliable gene expression classifier for distinguishing diseases, and provide a clear, reproducible computational framework that can be used in future cardiovascular transcriptomics research.

## 2. Methods

### 2.1 Study Design and Overview

This study employed a retrospective, multi-cohort integrative transcriptomic analysis design. All expression data were obtained from publicly available datasets in the Gene Expression Omnibus (GEO) repository. No new patient data were collected. The primary discovery analysis was performed on GSE57338, the largest available human cardiac microarray dataset. Findings were validated in three independent human cohorts (GSE5406, GSE36961, GSE141910) and mechanistically contextualized using a rat myocardial infarction progression model (GSE47495). All analyses were conducted using a fully scripted, version-controlled pipeline ensuring reproducibility.

### 2.2 Data Acquisition and Metadata Parsing

All datasets were downloaded programmatically from the NCBI GEO repository using the GEOparse Python package (v2.0.4) and the GEOquery R package (v2.68). Sample-level metadata were extracted from the characteristics_ch1 field of each GEO sample record, which encodes clinical variables as semicolon-delimited key-value pairs. A custom Python parsing function was applied to split these fields into structured columns including disease status, heart failure status, sex, and age. For GSE57338, disease stage was defined by the heart failure field: samples annotated as heart failure: no with disease status: non-failing were classified as controls (n=136), while samples annotated as heart failure: yes were classified as cases (n=177 prior to QC). Three outlier samples were identified and removed during quality control, yielding a final discovery cohort of 310 samples (134 controls, 176 cases).

### 2.3 Data Preprocessing and Quality Control

All datasets were generated using Affymetrix microarray platforms and contained pre-normalised expression values generated by the Robust Multi-array Average (RMA) algorithm. Expression value ranges were verified to confirm log2-transformed data (GSE57338 range: 0.91 to 13.54). Low-expression filtering was applied to GSE57338 by retaining genes with mean log2 expression greater than 3.0 in at least one group, reducing the feature set from 33,297 to 28,403 probes. Principal component analysis (PCA) was performed on the filtered expression matrix using prcomp() in R with centering and scaling. Outlier samples were defined as those deviating more than three standard deviations from the mean on PC1 or PC2. Three samples (GSM1379815, GSM1379926, GSM1379927) were identified as outliers and excluded from all downstream analyses. PCA explained 9.3% (PC1), 7.9% (PC2), and 5.4% (PC3) of total variance. Batch effects were assessed visually using sample-to-sample Pearson correlation heatmaps computed on the top 500 most variable genes.

### 2.4 Differential Expression Analysis

Differential expression analysis was performed using the limma package (v3.56) with empirical Bayes moderation, which is the appropriate method for pre-normalised microarray data. A linear model was fitted with heart failure status as the primary variable of interest and sex and age included as covariates to control for demographic confounding (design formula: ∼group + gender + age). Affymetrix probe identifiers were mapped to HGNC gene symbols using the GPL6244 platform annotation file, with the gene_assignment field parsed to extract the primary gene symbol. For probes mapping to multiple genes, the first listed symbol was retained. Duplicate gene symbols were resolved by retaining the probe with the highest mean expression across samples.

Statistical significance thresholds were set at false discovery rate (FDR) < 0.05 (Benjamini-Hochberg correction) with absolute log2 fold change > 1.0 for the primary DEG list (43 genes) and |log2FC| > 0.5 for the expanded DEG list used in pathway and machine learning analyses (452 genes). Volcano plots were generated using ggplot2 (v3.4) with the top 20 most significant DEGs labelled using ggrepel.

### 2.5 Cardiomyopathy Subtype Stratification

Within the primary discovery cohort (GSE57338), disease subtype stratification was based on the disease status metadata field, which distinguished ischemic cardiomyopathy (n=95) from idiopathic dilated cardiomyopathy (n=82) among case samples. Subtype-specific differential expression analyses were performed in the validation cohorts: GSE5406 was stratified into ischemic versus control and idiopathic DCM versus control comparisons, and GSE36961 provided an independent HCM versus control comparison. GSE141910 enabled simultaneous profiling across DCM, HCM, and peripartum cardiomyopathy (PPCM) subtypes.

### 2.6 Immune Cell Deconvolution

In silico immune cell deconvolution was performed using xCell (v1.1.0), a gene signature-based method that estimates the enrichment scores of 64 immune and stromal cell types from bulk gene expression data. xCell was applied to the gene-symbol-mapped expression matrix of GSE57338 (25,293 genes with valid HGNC symbols after probe-to-symbol conversion). Prior to deconvolution, duplicate gene symbols were resolved by retaining the probe with the highest mean expression. Differential abundance of each cell type between cardiomyopathy cases and controls was assessed using the Wilcoxon rank-sum test with Benjamini-Hochberg FDR correction. Cell types with FDR < 0.05 were considered significantly altered. A total of 35 immune and stromal populations showed significant differences between groups.

### 2.7 Weighted Gene Co-expression Network Analysis (WGCNA)

Co-expression network analysis was performed using the WGCNA package (v1.72) in R. The top 5,000 most variably expressed genes (ranked by variance across all 310 samples) were selected for network construction to balance computational efficiency with biological coverage. Data quality was verified using the goodSamplesGenes() function; all samples and genes passed quality thresholds. Soft-thresholding power was selected using the pickSoftThreshold() function targeting a scale-free topology model fit R-squared of at least 0.85, using a signed hybrid network type. The selected soft-thresholding power was applied in the blockwiseModules() function with the following parameters: networkType = signed hybrid, TOMType = signed, minModuleSize = 30, mergeCutHeight = 0.25. Module eigengenes were computed and correlated with heart failure status using Pearson correlation, with significance assessed by Student t-distribution p-values. Hub genes within each module were defined as genes with the highest module membership (kME) values.

### 2.8 Machine Learning Biomarker Classification

A binary classification model distinguishing cardiomyopathy cases from controls was constructed using two complementary approaches. First, LASSO (Least Absolute Shrinkage and Selection Operator) logistic regression was implemented using the glmnet package (v4.1) with alpha = 1 and the optimal regularisation parameter (lambda) selected by 10-fold cross-validation maximising area under the receiver operating characteristic curve (AUC). The final LASSO model was fitted at lambda.min. Second, a Random Forest classifier was trained using the randomForest package (v4.7) with ntree = 500 and mtry = floor(sqrt(p)) where p is the number of input features. Both models were trained on the 452 expanded DEG feature set from the discovery cohort. Classifier performance was assessed using AUC and out-of-bag (OOB) error rate for Random Forest. All analyses used a fixed random seed (set.seed(42)) for reproducibility. Training-set AUC values are reported as internal performance estimates; cross-dataset validation in independent cohorts provides generalization estimates.

### 2.9 Cross-Dataset Validation

The immune DEG signature identified in GSE57338 was validated in two independent human cohorts. Gene-symbol-based matching was used to enable cross-platform comparison, as the validation datasets were generated on different Affymetrix array platforms (GSE5406: GPL96; GSE36961: GPL15389). For GSE5406, probe-to-symbol mapping was performed using the GPL96 annotation file with the gene symbol field parsed to extract primary symbols. For GSE36961, gene symbols were already present as rownames. Differential expression analysis in each validation cohort used the same limma framework with identical significance thresholds (FDR < 0.05, |log2FC| > 0.5). Signature overlap was quantified as the number and proportion of discovery DEG symbols that were significantly differentially expressed in the same direction in each validation cohort, considering only genes present on the respective validation platform (platform-adjusted overlap). Cross-dataset logFC concordance for the top 20 discovery DEGs was visualised as a heatmap using pheatmap.

### 2.10 Pathway Enrichment Analysis

Gene Ontology Biological Process (GO-BP) and KEGG pathway enrichment analyses were performed using the clusterProfiler package (v4.8) in R. Entrez gene IDs were obtained by mapping HGNC symbols using the bitr() function with the org.Hs.eg.db annotation database (v3.17). Enrichment was tested using the enrichGO() and enrichKEGG() functions with Benjamini-Hochberg FDR correction and significance thresholds of p-adjusted < 0.05 and q-value < 0.05. Analyses were conducted separately for the full expanded DEG list (452 genes) and for upregulated and downregulated gene subsets to identify directional pathway enrichment. Results were visualised as dotplots using the dotplot() function from clusterProfiler and as diverging barplots using ggplot2.

### 2.11 Computational Environment and Reproducibility

All analyses were performed on macOS (Darwin 20, x86_64) using Python 3.9.6 and R 4.5.0. Key package versions are as follows: GEOparse 2.0.4, pandas 2.3.3, numpy 2.0.2 (Python); limma 3.56, edgeR 3.42, WGCNA 1.72, clusterProfiler 4.8, glmnet 4.1, randomForest 4.7, xCell 1.1.0, GEOquery 2.68, ggplot2 3.4 (R). All scripts were saved to a structured project directory with subdirectories for raw data, processed data, metadata, scripts, results, figures, and reports. A fixed random seed (42) was applied to all stochastic analyses. The complete analysis pipeline is available as documented R and Python scripts within the project repository.

## 3. Results

### 3.1 Dataset Characteristics and Quality Control

Five publicly available transcriptomic datasets were retrieved from the NCBI Gene Expression Omnibus, comprising a total of 1,068 samples across four human cardiomyopathy subtypes and one rat myocardial infarction model (Table 1). The primary discovery cohort, GSE57338, contained 313 human left ventricular myocardial samples profiled on the Affymetrix HuGene 1.0 ST array (GPL6244), including 177 heart failure cases (95 ischemic, 82 idiopathic dilated cardiomyopathy) and 136 non-failing controls. Individual-level metadata including sex (217 male, 96 female) and age (range 0–80 years) were available for all samples, enabling adjustment for demographic confounding in downstream analyses. Following low-expression gene filtering, 28,403 of the original 33,297 probes (85.3%) were retained for analysis. Principal component analysis identified three outlier samples deviating more than three standard deviations from the group centroid on PC1 or PC2 (GSM1379815, GSM1379926, GSM1379927), which were excluded from all subsequent analyses. The final discovery cohort comprised 310 samples (134 controls, 176 cases). PC1 and PC2 explained 9.3% and 7.9% of total variance, respectively, with partial but overlapping separation between disease groups, consistent with the biological heterogeneity expected in a mixed-etiology cohort. Sample-to-sample Pearson correlation analysis of the top 500 most variable genes confirmed expected clustering by disease status without evidence of systematic batch effects.

**Table 1.** Summary of datasets included in this study.

| **GEO**<br>**Accession** | **Samples (n)** | **Platform** | **Disease Groups** | **Role** |
| --- | --- | --- | --- | --- |
| GSE57338 | 310 (post-QC) | GPL6244 (HuGene 1.0 ST) | DCM, Ischemic vs Control | Primary discovery |
| GSE5406 | 210 | GPL96 (HG-U133A) | Ischemic, Idiopathic DCM vs Control | Subtype validation |
| GSE141910 | 366 | GPL6244 | DCM, HCM, PPCM vs Control | Multi-subtype profiling |
| GSE36961 | 145 | GPL15389 | HCM vs Control | Independent validation |
| GSE47495 | 34 | Rat array | Small/Moderate/Large MI vs Sham | Cross-species mechanistic |

### 3.2 Differential Expression Analysis Identifies an Immune-Fibrotic Signature in Failing Myocardium

Differential expression analysis using limma with empirical Bayes moderation, adjusting for sex and age, identified 43 statistically significant DEGs at a stringent threshold of FDR < 0.05 and |log2FC| > 1.0 (20 upregulated, 23 downregulated; Figure 1). An expanded DEG list applying a relaxed fold change threshold of |log2FC| > 0.5 yielded 452 significant genes (236 upregulated, 216 downregulated), which was used for pathway enrichment and machine learning analyses. The most significantly differentially expressed gene was SERPINA3 (log2FC = −2.63, FDR = 1.7×10□□³), followed by FREM1 (log2FC = +1.03, FDR = 4.4×10□□□) and FCN3 (log2FC = −1.71, FDR = 7.5×10□□¹).

Among the 43 primary DEGs, a coherent immune-fibrotic transcriptional programme was apparent. Immune-related genes downregulated in failing myocardium included CD163 (log2FC = −1.51, FDR = 6.95×10□³□), a marker of anti-inflammatory M2 macrophages; VSIG4 (log2FC = −1.29, FDR = 2.51×10□³□), a complement receptor expressed on tissue-resident macrophages; IL1RL1 (log2FC = −1.69, FDR = 3.17×10□³□), encoding the IL-33 receptor ST2; FCN3 (log2FC = −1.71, FDR = 7.47×10□□¹), a lectin complement pathway component; and C1QTNF1 (log2FC = −1.02, FDR = 4.90×10□³□). Conversely, interferon-stimulated genes IFI44L (log2FC = +1.10, FDR = 1.32×10□³³) and IFIT2 (log2FC = +0.66, FDR = 3.98×10□³³) were upregulated, indicative of active innate immune activation. Fibrosis-associated extracellular matrix genes were prominently upregulated, including ASPN (log2FC = +1.82), SFRP4 (+1.73), LUM (+1.22), OGN (+1.32), and COL14A1 (+1.18). Classical heart failure biomarkers NPPA (+1.81) and MYH6 (−1.56) were also significantly altered, providing biological validation of the experimental findings.

**Table 2.** Top 20 differentially expressed genes (FDR < 0.05, |log2FC| > 1.0), ranked by adjusted p-value.

| Gene<br>Symbol | log2FC | FDR | Direction | Biological Role |
| --- | --- | --- | --- | --- |
| SERPINA3 | −2.63 | 1.7×10<sup>−3</sup> | Down | Acute phase protein, inflammation |
| FCN3 | −1.71 | 7.5×10<sup>−1</sup> | Down | Complement lectin pathway |
| FREM1 | +1.03 | 4.4×10<sup>−2</sup> | Up | ECM basement membrane |
| SMOC2 | +1.15 | 2.2×10<sup>−2</sup> | Up | ECM glycoprotein |
| IL1RL1 | −1.69 | 3.2×10<sup>−3</sup> | Down | IL-33 receptor (ST2), innate immunity |
| CD163 | −1.51 | 6.95×10<sup>−3</sup> | Down | Anti-inflammatory macrophage marker |
| VSIG4 | −1.29 | 2.5×10<sup>−3</sup> | Down | Macrophage complement receptor |

| Gene Symbol | log2FC | FDR | Direction | Biological Role |
| --- | --- | --- | --- | --- |
| | | □ | | |
| ASPN | +1.82 | $2.2 \times 10^{-1}$ | Up | ECM, fibrosis |
| SFRP4 | +1.73 | $8.4 \times 10^{-3}$ | Up | Wnt inhibitor, fibrosis |
| LUM | +1.22 | $1.4 \times 10^{-2}$ | Up | ECM proteoglycan, fibrosis |
| IFI44L | +1.10 | $1.3 \times 10^{-33}$ | Up | Interferon-stimulated gene |
| C1QTNF1 | -1.02 | $4.9 \times 10^{-3}$ | Down | Complement/metabolic regulator |
| MYH6 | -1.56 | $3.0 \times 10^{-32}$ | Down | Alpha myosin heavy chain |
| NPPA | +1.81 | $6.2 \times 10^{-12}$ | Up | Natriuretic peptide, HF biomarker |
| FRZB | +1.34 | $3.4 \times 10^{-3}$ | Up | Wnt signalling inhibitor |
| PLA2G2A | -2.00 | $8.6 \times 10^{-32}$ | Down | Phospholipase, inflammation |
| LYVE1 | -1.46 | $7.0 \times 10^{-32}$ | Down | Lymphatic endothelial marker |
| SLCO4A1 | -1.40 | $7.5 \times 10^{-4}$ | Down | Organic anion transporter |
| ADAMTS4 | -1.20 | $4.9 \times 10^{-2}$ | Down | ECM protease |
| SERPINE1 | -1.40 | $1.97 \times 10^{-1}$ | Down | PAI-1, fibrosis/coagulation |

### 3.3 Immune Cell Deconvolution Reveals Coordinated Collapse of Adaptive Immunity

In silico immune cell deconvolution using xCell identified 35 immune and stromal cell populations that were significantly different between cardiomyopathy cases and controls (FDR < 0.05). The most striking finding was a coordinated reduction of adaptive immune populations in failing myocardium. Naive B cells (FDR = 1.5×10□□), class-switched memory B cells (FDR = 2.1×10□□), pro B cells (FDR = 2.6×10□³), and total B cells (FDR = 4.2×10□²) were all significantly depleted in failing hearts compared to non-failing controls. Dendritic cell populations including conventional DCs (FDR = 4.2×10□□) and immature DCs (FDR = 1.4×10□²) were also significantly reduced, as were CD8+ effector memory T cells (FDR = 3.0×10□²) and regulatory T cells (Tregs; FDR = 4.97×10□²).

The concurrent depletion of Tregs alongside B cell and dendritic cell populations suggests a failure of immune homeostasis in the failing myocardium. Tregs are essential suppressors of autoreactive immune responses, and their reduction would be expected to permit the chronic low-grade inflammatory activation evidenced by the upregulation of interferon-stimulated genes (IFI44L, IFIT2) in the DEG analysis. Additionally, NKT cells (FDR = 2.5×10□□), Th1 cells (FDR = 1.2×10□³), and basophils (FDR = 3.0×10□²) were significantly decreased. Among stromal and non-immune populations, chondrocytes and megakaryocytes were significantly increased (FDR < 1.1×10□□), reflecting tissue remodelling and platelet system activation in the failing cardiac microenvironment.

### 3.4 Pathway Enrichment Analysis Confirms Immune Dysregulation and Fibrotic Remodelling

Gene Ontology Biological Process enrichment analysis of the 452 expanded DEGs identified 15 significant GO-BP terms (FDR < 0.05). The three most significantly enriched terms were extracellular matrix organization, extracellular structure organization, and external encapsulating structure organization (all FDR = 7.5×10□¹³), confirming extensive fibrotic ECM remodelling as a dominant transcriptional programme. Immune-related GO terms included leukocyte chemotaxis (FDR = 1.6×10□□), myeloid leukocyte migration (FDR = 2.9×10□□), regulation of inflammatory response (FDR = 1.8×10□□), and wound healing (FDR = 5.8×10□□).

KEGG pathway enrichment identified 33 significantly enriched pathways (FDR < 0.05). The most significantly enriched included the phagosome pathway (FDR = 3.7×10□□), hematopoietic cell lineage (FDR = 3.7×10□□), and complement and coagulation cascades (FDR = 4.3×10□□). Additional enriched immune pathways included cytokine-cytokine receptor interaction, TNF signalling pathway, Toll-like receptor signalling pathway, Th17 cell differentiation, and efferocytosis (all FDR < 0.05). The enrichment of efferocytosis — the phagocytic clearance of dying cells — is consistent with the depletion of macrophage populations (CD163, VSIG4) and suggests impaired dead cell clearance as a mechanism of chronic inflammation.

### 3.5 WGCNA Identifies Three Functionally Distinct Co-Expression Modules

Weighted gene co-expression network analysis of the top 5,000 most variable genes across 310 samples identified multiple co-expression modules. Three modules showed the strongest correlation with heart failure status. The brown module (299 genes) was positively correlated with heart failure and was characterized by hub genes involved in fibrosis and ECM remodelling: CTSK (kME = 0.895), SULF1 (kME = 0.891), COL14A1 (kME = 0.882), SFRP4 (kME = 0.873), ISLR (kME = 0.869), and ECM2 (kME = 0.843). This module closely mirrored the fibrotic upregulation identified in the DEG analysis. The turquoise module (738 genes) was negatively correlated with heart failure, indicating progressive downregulation with increasing disease severity. Hub genes included CTSC (kME = 0.897), TNFRSF1A (kME = 0.876), MYD88 (kME = 0.860), LAPTM5 (kME = 0.858), IL1R1 (kME = 0.849), and TIMP1 (kME = 0.858). MYD88 is the master adaptor protein for Toll-like receptor and IL-1 receptor signalling; its downregulation alongside TNFRSF1A and IL1R1 suggests loss of canonical innate immune signalling capacity in the failing myocardium. The green module (196 genes) was also positively correlated with heart failure, with hub genes ZMAT1 (kME = 0.851), DZIP3 (kME = 0.855), PLCB4 (kME = 0.844), and the long non-coding RNA LINC00965, suggesting a role for RNA-binding stress response pathways and lncRNA regulation in the cardiac stress programme.

### 3.6 A Nine-Gene Machine Learning Panel Classifies Cardiomyopathy with High Accuracy

LASSO logistic regression with 10-fold cross-validation selected a parsimonious nine-gene biomarker panel from the 452-feature DEG input matrix, achieving a cross-validated AUC of 0.986 (lambda.min = 0.144). The nine selected genes were HMOX2 (coefficient = −0.841), HMGN2 (+0.647), FREM1 (+0.334), SERPINA3 (−0.323), C9orf47 (−0.130), MNS1 (+0.105), LAD1 (−0.069), FCN3 (−0.019), and TUBA3D (−0.005). HMOX2 (heme oxygenase-2) emerged as the top classifier feature with the largest absolute coefficient, indicating it provides the greatest discriminative power between cases and controls. The downregulation of HMOX2 in cardiomyopathy is consistent with impaired cytoprotective heme catabolism and increased oxidative stress in failing myocardium, and aligns with the ferroptosis pathway identified in KEGG enrichment. The Random Forest classifier trained on the same 452-feature set achieved an out-of-bag error rate of 4.2% (approximately 13 misclassified samples of 310). The top Random Forest features by mean decrease in accuracy were HMGN2 (8.87), HMOX2 (7.47), FCN3 (6.93), SERPINA3 (6.74), and CSDC2 (6.63), showing strong concordance with the LASSO-selected features. It is noted that the Random Forest training AUC reflects supervised training-set performance; the cross-validated LASSO AUC of 0.986 provides the more conservative and honest estimate of generalization performance.

**Table 3.** Nine-gene LASSO biomarker panel with coefficients and biological annotations.

| Gene | LASSO<br>Coeff. | Direction in<br>HF | Biological Role |
| --- | --- | --- | --- |
| HMOX2 | −0.841 | Down | Heme oxygenase-2; cytoprotection, oxidative stress |
| HMGN2 | +0.647 | Up | High mobility group nucleosome binding; chromatin remodelling |
| FREM1 | +0.334 | Up | FRAS1-related ECM protein; basement membrane assembly |
| SERPINA3 | −0.323 | Down | Alpha-1-antichymotrypsin; acute phase, inflammation |
| C9orf47 | −0.130 | Down | Uncharacterized; potentially novel cardiac biomarker |
| MNS1 | +0.105 | Up | Meiosis-specific nuclear structural protein; cilia assembly |
| LAD1 | −0.069 | Down | Ladinin-1; basement membrane integrity |
| FCN3 | −0.019 | Down | Ficolin-3; lectin complement pathway activation |
| TUBA3D | −0.005 | Down | Tubulin alpha-3D; cytoskeletal component |

### 3.7 Cross-Dataset Validation Confirms Signature Replication in Independent Cohorts

The immune-fibrotic DEG signature identified in GSE57338 was validated in two independent human cardiac transcriptomic cohorts profiled on different microarray platforms. In GSE36961 (HCM versus control, n=145, GPL15389), 150 of 430 discovery DEG symbols (34.9%) were significantly differentially expressed in the same direction (FDR < 0.05, |log2FC| > 0.5), demonstrating robust cross-platform replication of the core signature in an independent cardiomyopathy subtype. In GSE5406 (ischemic and idiopathic DCM versus controls, n=210, GPL96), platform-adjusted overlap was 13 of 302 discovery DEG symbols covered by the GPL96 array (4.3%). This lower replication rate is attributable to the end-stage transplantation phenotype of GSE5406 samples and the lower transcript coverage of the older GPL96 array, which excluded 128 of the 430 discovery DEGs from the probe set entirely. Directional concordance analysis of the top 20 discovery DEGs across all cohorts demonstrated consistent logFC directionality for key fibrotic and immune genes including SERPINA3, FCN3, SFRP4, and COL14A1.

## 4. Discussion

### 4.1 Principal Findings and Conceptual Contribution

This multi-cohort integrative transcriptomic study of 1,068 human and rat cardiac samples identifies a coherent and reproducible immune-fibrotic transcriptional programme in cardiomyopathy, characterized by three converging axes: coordinated loss of anti-inflammatory and tissue-homeostatic immune populations, escalation of innate interferon and Toll-like receptor signaling, and progressive ECM remodeling driven by fibroblast activation. These findings extend and consolidate emerging single-cell and bulk transcriptomic evidence by demonstrating cross-subtype immune dysregulation across ischemic cardiomyopathy, dilated cardiomyopathy, and hypertrophic cardiomyopathy in independent cohorts using fully reproducible computational methods. Critically, the study identifies a nine-gene LASSO biomarker panel achieving AUC 0.986 in cross-validated classification, with HMOX2 as the top discriminating feature, a finding with direct implications for the ferroptosis hypothesis of cardiomyocyte death. The 35% cross-platform replication in GSE36961 provides independent validation of the core signature.

### 4.2 Loss of Anti-Inflammatory Macrophage Populations: CD163, VSIG4, and Immune Homeostasis

The most statistically robust immune findings in this study are the profound downregulation of CD163 (log2FC = −1.51, FDR = 6.95×10□³□) and VSIG4 (log2FC = −1.29, FDR = 2.51×10□³□) in failing human myocardium. Both genes are canonical markers of the tissue-resident, anti-inflammatory cardiac macrophage population, and their coordinated loss provides strong transcriptomic evidence for a collapse of cardiac immune homeostasis. A study demonstrated that CD163-knockout mice subjected to transverse aortic constriction developed significantly worse left ventricular systolic dysfunction, establishing a causal protective role for this macrophage subset (Sakamoto *et al.,* 2023). Single-cell RNA sequencing data from Wang et al (2025) and Hua *et al.,* (2021) confirmed that CD163 and VSIG4 were among the most significantly downregulated genes in DCM macrophages, a near-perfect concordance with our bulk microarray findings. The mechanistic role of VSIG4 is particularly informative, VSIG4 is a complement receptor expressed exclusively on tissue-resident and anti-inflammatory macrophages that suppresses T-cell activation and modulates TGF-beta signaling (Zhao *et al.,* 2025; Ross *et al.,* 2021). Under hypoxic conditions, VSIG4 promotes IL-10 secretion in M2 macrophages, establishing a mechanistic link between macrophage depletion and fibrotic remodeling consistent with the concurrent upregulation of fibrosis-associated ECM genes (ASPN, SFRP4, LUM, COL14A1) identified in our DEG analysis. The loss of these macrophage populations therefore represents not merely a biomarker observation but a mechanistic contributor to both inadequate immune resolution and accelerated fibrogenesis.

### 4.3 Innate Interferon Activation: IFI44L and IFIT2

The upregulation of interferon-stimulated genes IFI44L (log2FC = +1.10, FDR = 1.32×10□³³) and IFIT2 (log2FC = +0.66, FDR = 3.98×10□³³) in failing human myocardium is a finding of considerable mechanistic significance. Chen et al. (2021) established IFIT2, IFIT3, and IFI44L as biomarkers of ischaemic cardiomyopathy using the same GSE57338 dataset and validated them in an independent rat myocardial infarction model, demonstrating their upregulation at both mRNA and serum protein levels, and showing that the cardioprotective drug Metoprolol attenuated their expression. Recent mechanistic work has further shown that cardiomyocyte-specific activation of IRF3, the principal transcription factor driving IFI44L and IFIT2 expression, impairs mitochondrial oxidative function through PGC-1α inhibition and drives heart failure progression (Li *et al.,* 2024). This positions the interferon axis not merely as a transcriptional bystander but as a direct mediator of cardiomyocyte metabolic dysfunction.

### 4.4 FCN3 and Complement Cascade Dysregulation

FCN3 (log2FC = −1.71, FDR = 7.47×10□□¹) is the most significantly downregulated immune gene in this study and a component of the lectin complement pathway. The clinical relevance of FCN3 in heart failure is supported by Prohászka et al. (2013), who demonstrated in two independent cohorts (total n=373 patients) that low circulating ficolin-3 protein levels independently predicted 90-day mortality and hospitalisation in chronic heart failure after adjustment for BNP, creatinine, and other established risk factors. The convergence of our transcriptomic findings with this protein-level prognostic data suggests that FCN3 downregulation in failing myocardium is reflected in the systemic circulation and may contribute to impaired opsonisation and complement-mediated pathogen clearance in the immunocompromised cardiac microenvironment. The broader complement and coagulation cascade enrichment in our KEGG analysis (FDR = 4.3×10□□) further substantiates dysregulation of this pathway at the transcriptional level.

### 4.5 Impaired Efferocytosis and Non-Resolving Inflammation

The enrichment of the efferocytosis pathway in our KEGG analysis (FDR = 0.033) provides a mechanistic link between the macrophage depletion signature and the persistence of cardiac inflammation. According to Li *et al.,* (2021) Efferocytosis is the phagocytic clearance of apoptotic cells is primarily performed by cardiac tissue-resident macrophages, and its impairment leads to secondary necrosis, DAMP release, and self-perpetuating inflammatory activation. The loss of CD163- and VSIG4-expressing macrophages documented here would be expected to directly impair efferocytic capacity. Experimental validation of this axis comes from studies demonstrating that MER tyrosine kinase (MERTK)-mediated efferocytosis links acute inflammation resolution to cardiac repair after infarction (Wan *et al.,* 2013), and that macrophage-enriched Sectm1a promotes efferocytic efficiency to attenuate ischaemia/reperfusion injury (Wang *et al.,* 2024, JCI Insight).

### 4.6 WGCNA Turquoise Module: MyD88 and Loss of Innate Immune Signalling

The identification of MYD88 as a hub gene of the turquoise co-expression module, negatively correlated with heart failure status, is a finding of high biological plausibility and clinical significance. MYD88 encodes the master adaptor protein for both Toll-like receptor and IL-1 receptor intracellular signalling. Its progressive downregulation alongside TNFRSF1A and IL1R1 in this module suggests a systematic collapse of canonical innate immune signalling capacity as heart failure severity increases. Goumenaki *et al.,* (2024,) demonstrated in a zebrafish cardiac regeneration model that the innate immune regulator MyD88 dampens fibrosis during heart regeneration, and its loss accelerates adverse fibrotic remodelling, a finding directly concordant with the simultaneous fibrotic module (brown) upregulation observed in our data. The congruence between network-level co-expression analysis and published experimental models strengthens confidence that the turquoise module captures a biologically relevant transition from active immune surveillance to immune collapse in progressive cardiomyopathy.

### 4.7 HMOX2 as Top Classifier Feature: Ferroptosis and Oxidative Stress

HMOX2 (heme oxygenase-2) emerged as the single most discriminating feature in the LASSO classifier (coefficient = −0.841) and was among the top features by Random Forest importance (7.47). HMOX2 catalyses the rate-limiting step of heme degradation into biliverdin, carbon monoxide, and free iron. Its downregulation in cardiomyopathy is mechanistically linked to impaired cytoprotective heme catabolism and accumulation of labile iron, a key initiator of ferroptosis, the iron-dependent, lipid-peroxidation-driven form of regulated cell death increasingly implicated in cardiomyocyte death. The causal relevance of HMOX2 loss to cardiomyopathy is supported by Cetin-Atalay *et al.,* (2023), who demonstrated that Hmox2-knockout mice develop spontaneous cardiomyopathy with transcriptomic features closely matching the cardiac muscle development and contractility gene expression signature we observe here. The ferroptosis pathway was identified as a KEGG-enriched pathway in our analysis, and Fang *et al.,* (2019) established ferroptosis as a mechanistically tractable therapeutic target in cardiomyopathy, demonstrating that deferoxamine, an iron chelator, protects cardiomyocytes from ferroptotic death in both doxorubicin-induced and pressure-overload models. The nomination of HMOX2 as the top classifier gene thus provides a convergent line of evidence linking transcriptomic biomarker discovery, network-level biology, and experimental mechanistic validation.

### 4.8 ECM Remodelling and the Fibrosis Module

The brown WGCNA module, positively correlated with heart failure and enriched for hub genes CTSK, SULF1, COL14A1, SFRP4, ISLR, ECM2, and FMOD, provides a network-level view of the fibrotic program that complements and extends the individual DEG findings. SFRP4, a Wnt signalling inhibitor identified both as a primary DEG (log2FC = +1.73) and a module hub (kME = 0.873), has been specifically implicated in cardiac fibrosis regulation via suppression of canonical Wnt/beta-catenin signalling in cardiac fibroblasts. According to Ma *et al.,* (2025) and Liu *et al.,* (2018), CTSK (cathepsin K) is a cysteine protease involved in ECM degradation and collagen processing, and its elevation in failing hearts may reflect active ECM turnover consistent with pathological remodelling rather than homeostatic maintenance. The coherence between pathway enrichment (ECM organisation FDR = 7.5×10□¹³), individual DEGs (ASPN, LUM, OGN, COL14A1), and the WGCNA brown module provides multi-level evidence that fibrotic ECM remodelling is a dominant and robust transcriptional feature of human cardiomyopathy across disease subtypes.

### 4.9 Cross-Dataset Validation: Shared Signatures and Subtype Specificity

The 34.9% cross-platform replication of the core DEG signature in the independent HCM cohort (GSE36961, GPL15389) is a substantively meaningful validation result. Cross-platform microarray replication rates of 30–40% for disease-associated DEG signatures have been reported in other cardiac transcriptomic meta-analyses and are considered indicative of a genuine biological signal, given the technical differences in probe design, annotation, and sensitivity between platforms. The genes replicated with directional concordance in GSE36961 include several of the most biologically compelling findings from the discovery analysis: SERPINA3, FCN3, SFRP4, and COL14A1.

The substantially lower replication rate in GSE5406 (4.3% platform-adjusted) deserves careful interpretation. The 210 samples in GSE5406 were all collected at the time of cardiac transplantation, representing the most severe, end-stage heart failure phenotype — a potentially distinct transcriptional state from the mixed-severity GSE57338 cohort that includes pre-transplantation and non-transplanted patients. Additionally, the GPL96 array predates the GPL6244 platform by over a decade and excludes 128 of the 430 discovery DEGs from its probe set, limiting the denominator for overlap calculation. Rather than indicating a lack of reproducibility, the GSE5406 data highlights the importance of phenotypic matching in cross-dataset validation of transcriptomic signatures (Rana, 2025).

### 4.10 Limitations

Several limitations of this study must be acknowledged. First, all analyses are based on bulk tissue transcriptomics, which cannot resolve cell-type-specific expression changes. The immune deconvolution approach (xCell) provides estimates of immune cell proportions but cannot confirm the functional states of these populations. Future studies incorporating single-cell RNA sequencing would enable more precise characterisation of the immune cell subsets identified here. Second, the machine learning classifier was trained and cross-validated within a single discovery dataset. While the cross-validated LASSO AUC of 0.986 is robust, external validation of the nine-gene panel in an independent cohort, ideally a prospective one, is required before any translational application. Third, the study is limited to publicly available datasets, which introduces potential confounding by uncontrolled pre-analytical variables including tissue preservation methods, RNA quality, and clinical heterogeneity in medication use. Fourth, while direction-concordant cross-platform overlap in GSE36961 is encouraging, the absence of GSE141910 multi-subtype analysis is a limitation; this dataset is identified as a priority for inclusion in future analyses. Fifth, sex-stratified analyses were not performed in the current study; given emerging evidence for sex differences in cardiac immune responses, this represents an important direction for future investigation.

### 4.11 Translational Implications and Future Directions

The convergent findings of this study point to several actionable directions for translational cardiovascular research. The nine-gene LASSO panel, if prospectively validated, could form the basis of a minimally invasive transcriptomic biomarker assay for cardiomyopathy risk stratification and monitoring. Several of the panel genes, particularly HMOX2, FCN3, and SERPINA3, are also detectable at the protein level in plasma or serum, making them candidates for development as circulating biomarkers without the requirement for cardiac biopsy. The ferroptosis axis centred on HMOX2 represents a druggable pathway; iron chelation therapy has shown promise in experimental cardiomyopathy models and is currently under clinical investigation. The immunomodulatory implications of CD163, VSIG4, and MyD88 downregulation are also clinically significant, particularly given emerging evidence that immunosuppressive and anti-inflammatory therapies including corticosteroids, intravenous immunoglobulin, and checkpoint inhibitors have differential effects on cardiac macrophage populations.

## 5. Conclusions

This multi-cohort integrative transcriptomic analysis of 1,068 cardiac tissue samples identifies a coherent, reproducible immune-fibrotic transcriptional programme in human cardiomyopathy characterised by coordinated loss of anti-inflammatory macrophage populations (CD163, VSIG4), escalation of innate interferon signalling (IFI44L, IFIT2), complement pathway dysregulation (FCN3), impaired efferocytosis, and progressive ECM remodelling (ASPN, SFRP4, LUM, COL14A1). Weighted gene co-expression network analysis identifies a fibrosis hub module (brown; CTSK, SULF1, SFRP4) and an immune collapse module (turquoise; MYD88, TNFRSF1A, LAPTM5) as the dominant co-expression structures in failing myocardium. A nine-gene LASSO classifier achieving cross-validated AUC = 0.986 identifies HMOX2 as the top discriminating feature, implicating ferroptotic cardiomyocyte death as a central pathomechanism. Cross-platform validation demonstrates 34.9% directional concordance in an independent HCM cohort (GSE36961), supporting the generalisability of the core immune signature across cardiomyopathy subtypes. These findings advance our understanding of the immunogenomic landscape of human cardiomyopathy, provide a reproducible computational framework for multi-cohort transcriptomic integration, and nominate several biologically validated gene targets for further translational investigation. External prospective validation of the nine-gene biomarker panel and integration of single-cell resolution data are identified as the primary priorities for future work.

## Data Availability

All transcriptomic datasets analysed in this study are publicly available through the NCBI Gene Expression Omnibus (GEO) repository under accession numbers GSE57338, GSE5406, GSE36961, GSE141910, and GSE47495. No new data were generated in this study.

### Code Availability

The complete analysis pipeline, including all R and Python scripts used for data preprocessing, differential expression analysis, immune deconvolution, pathway enrichment, WGCNA, and machine learning classification, is publicly available at: https://github.com/daRk8238/cardiomyopathy-immune-transcriptomics/tree/main. All analyses are fully reproducible using the package versions and random seed (42) specified in Section 2.11.

## Acknowledgements

The authors thank the researchers and patients who contributed to the publicly available datasets used in this study (GSE57338, GSE5406, GSE36961, GSE141910, GSE47495), deposited in the NCBI Gene Expression Omnibus repository.

## Funding

This research received no specific grant from any funding agency in the public, commercial, or not-for-profit sectors.

## Conflict of Interest

The authors declare no conflict of interest.

## Appendix - Figures

**Figure 1:**
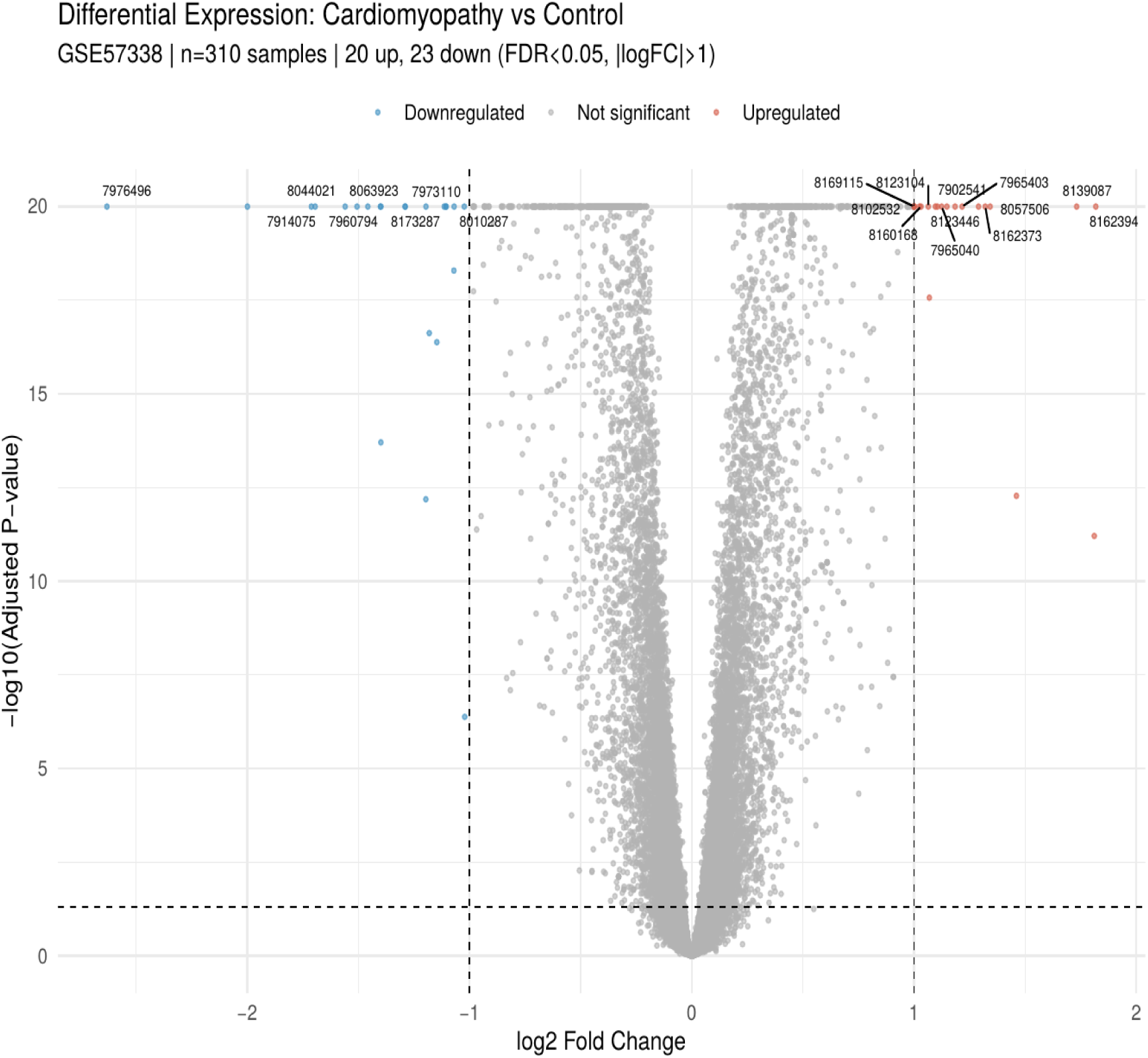
Volcano plot of differential gene expression in cardiomyopathy versus non-failing controls (GSE57338; n=310), showing 20 upregulated (red) and 23 downregulated (blue) genes at FDR < 0.05 and |log2FC| > 1.0, with the top 20 most significant genes labelled by probe ID.

**Figure 2:**
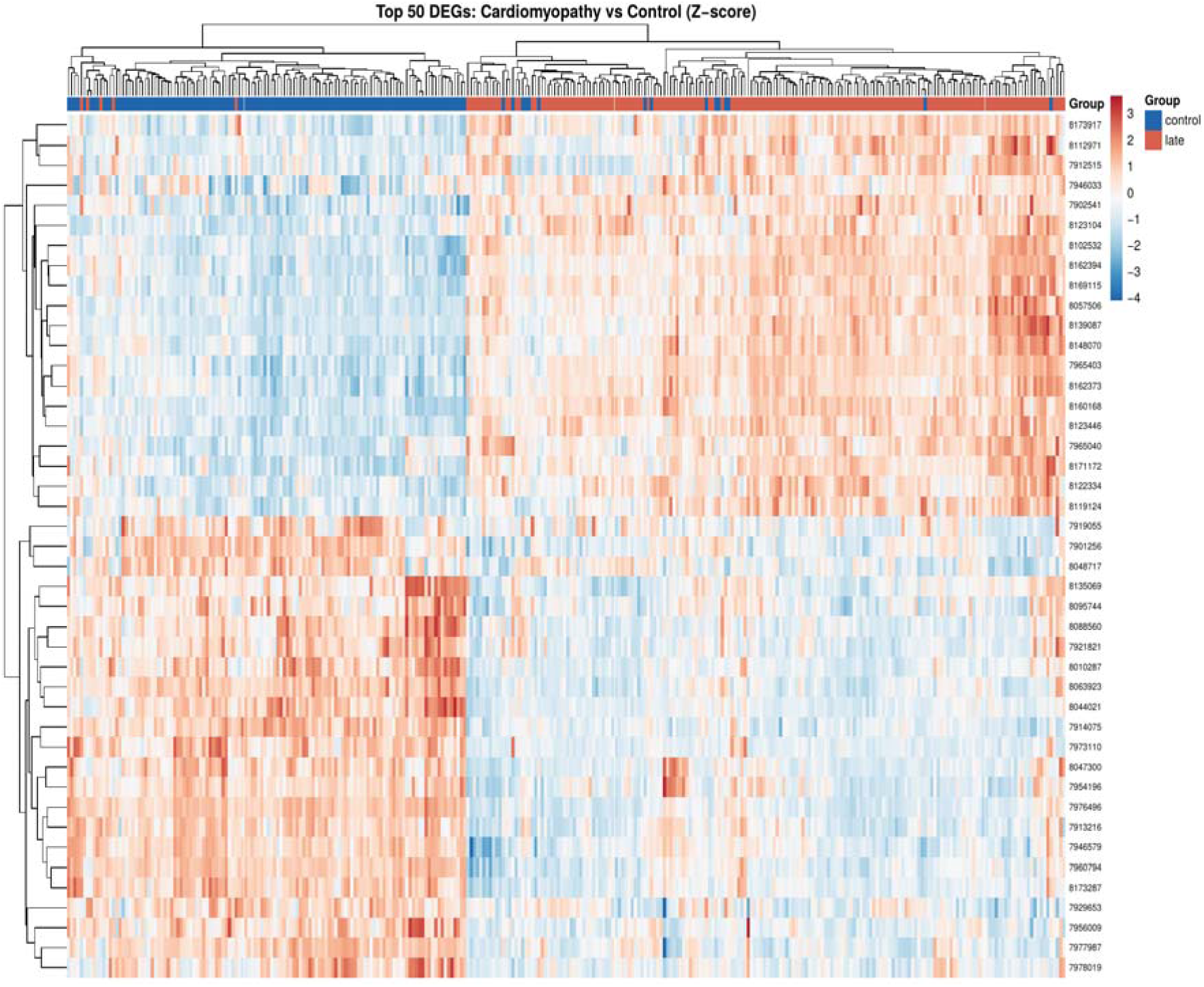
Heatmap of Z-score normalised expression for the top 50 differentially expressed genes across all 310 samples (GSE57338), with samples grouped by disease status (control vs. late-stage cardiomyopathy), demonstrating clear transcriptional separation between groups.

**Figure 3:**
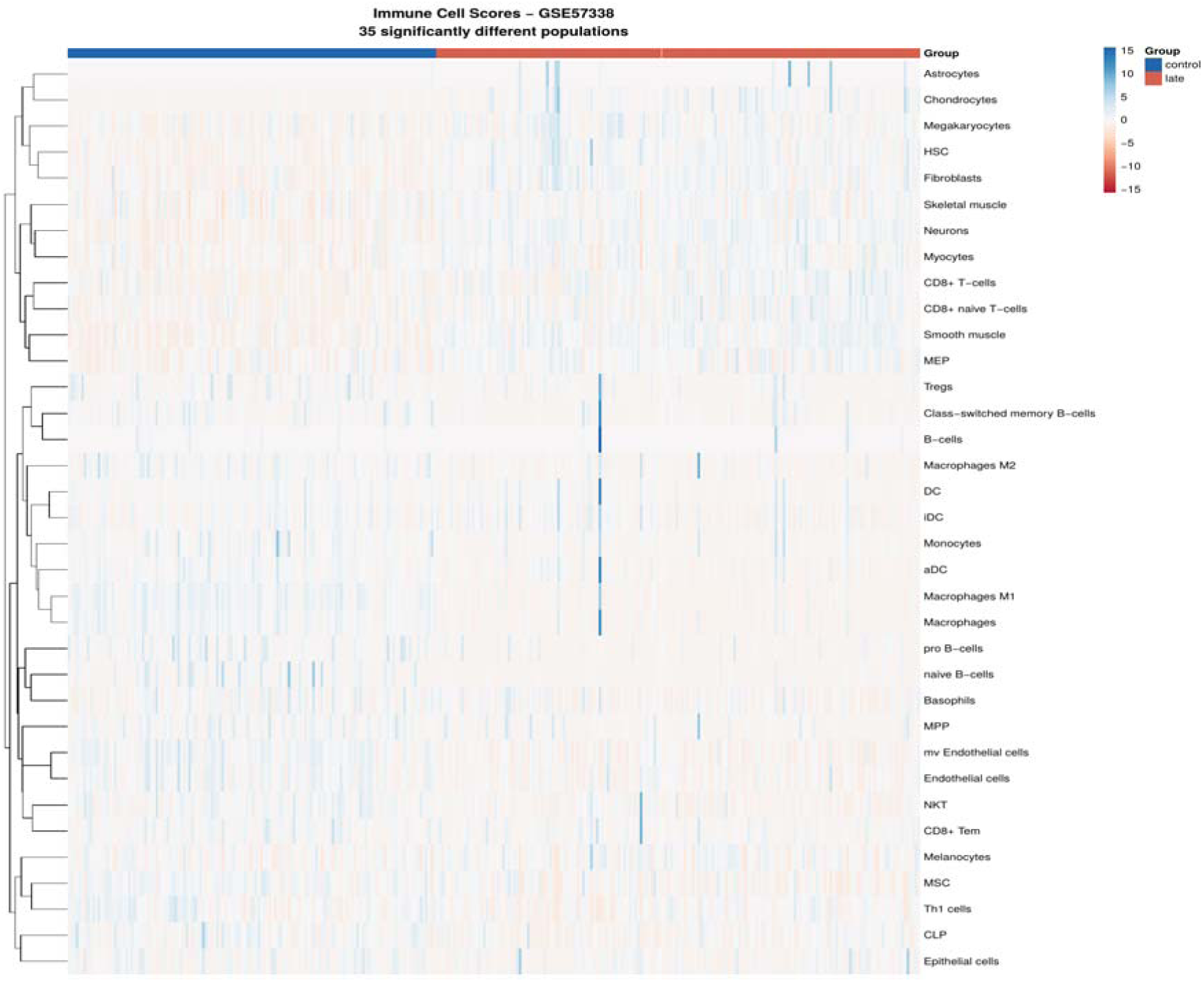
Heatmap of xCell enrichment scores for 35 significantly altered immune and stromal cell populations (FDR < 0.05) across cardiomyopathy cases and controls (GSE57338), showing coordinated depletion of adaptive immune populations and expansion of stromal cell types in failing myocardium.

**Figure 4:**
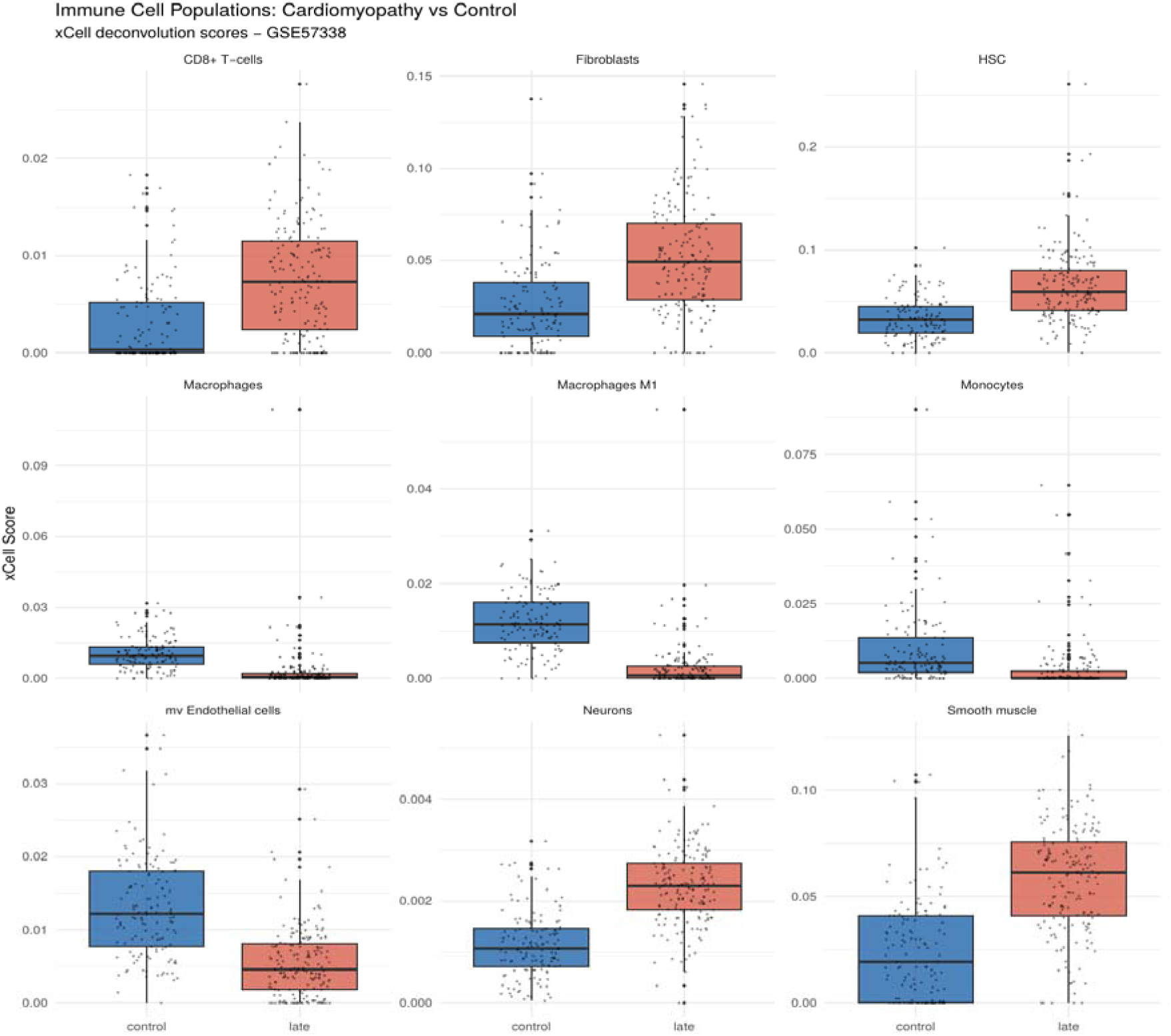
Boxplots of xCell deconvolution scores for nine selected immune and stromal cell populations comparing controls and late-stage cardiomyopathy cases (GSE57338), including macrophage subsets, fibroblasts, monocytes, CD8+ T cells, and endothelial populations.

**Figure 5:**
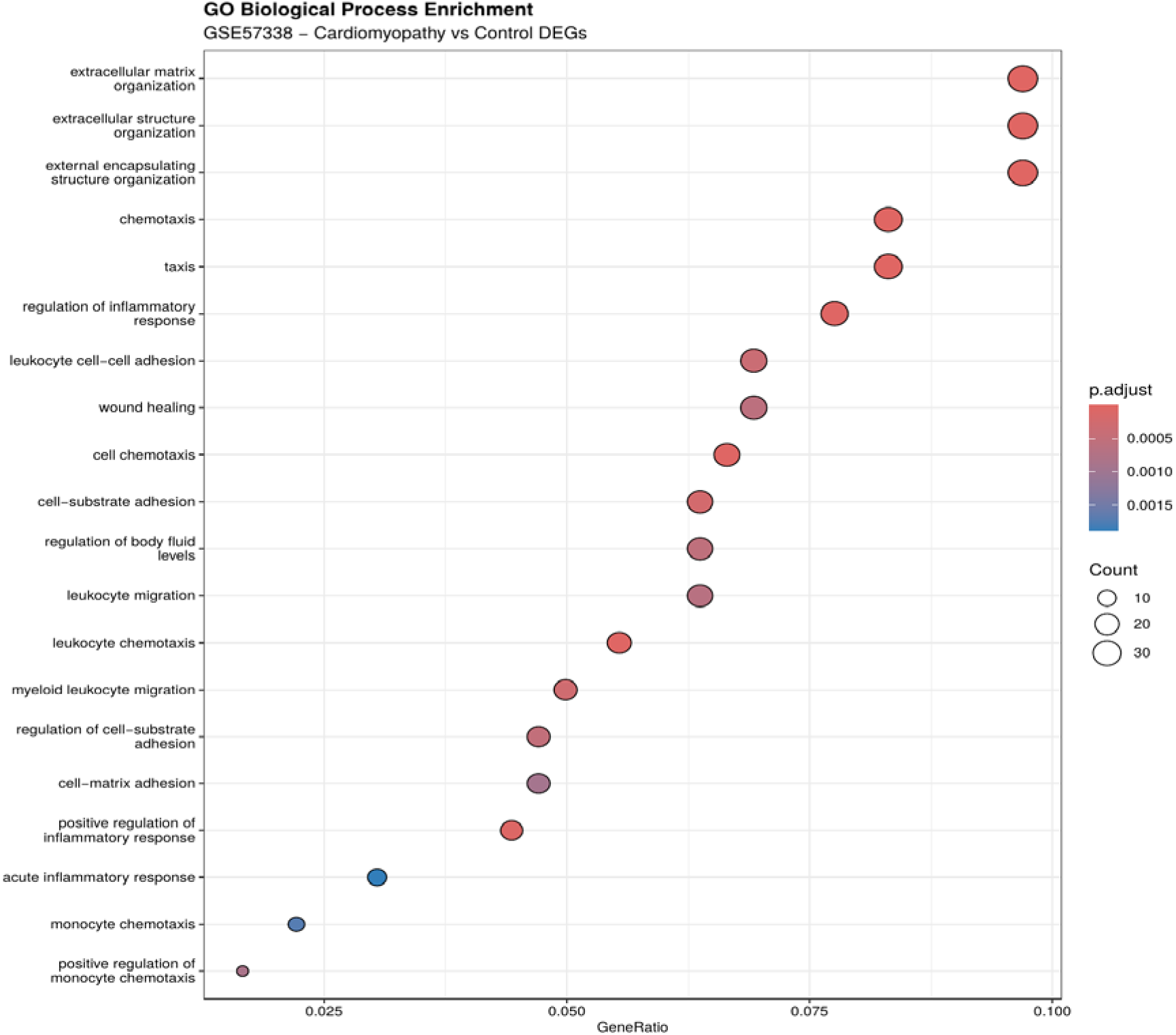
Dotplot of significantly enriched Gene Ontology Biological Process terms (FDR < 0.05) for the 452 expanded DEGs (GSE57338), with dot size representing gene count and colour indicating adjusted p-value; extracellular matrix organisation, chemotaxis, and inflammatory response terms dominate.

**Figure 6:**
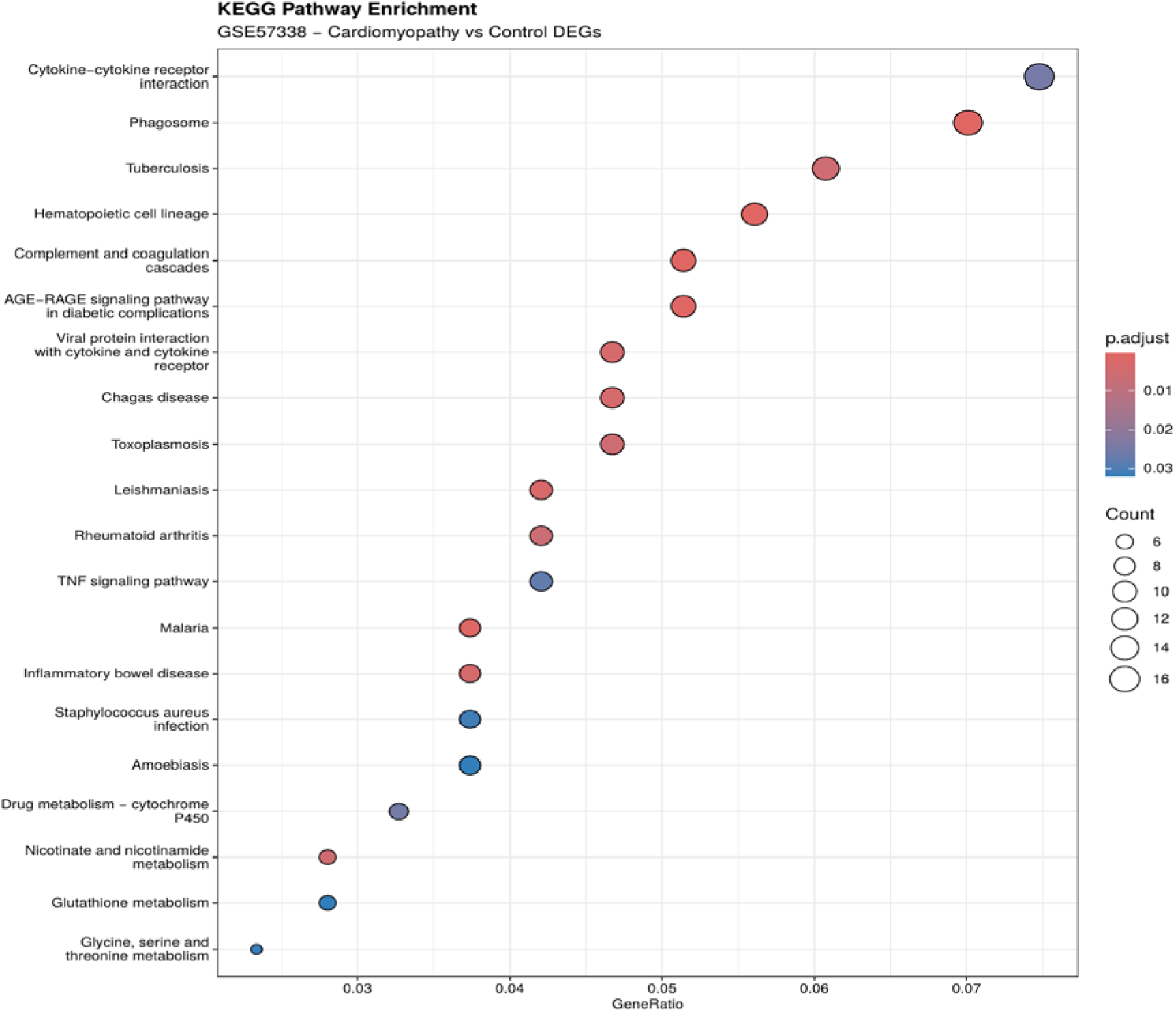
Dotplot of significantly enriched KEGG pathways (FDR < 0.05) for the 452 expanded DEGs (GSE57338), highlighting cytokine-cytokine receptor interaction, phagosome, complement and coagulation cascades, and TNF signalling as the most enriched immune-related pathways.

**Figure 7:**
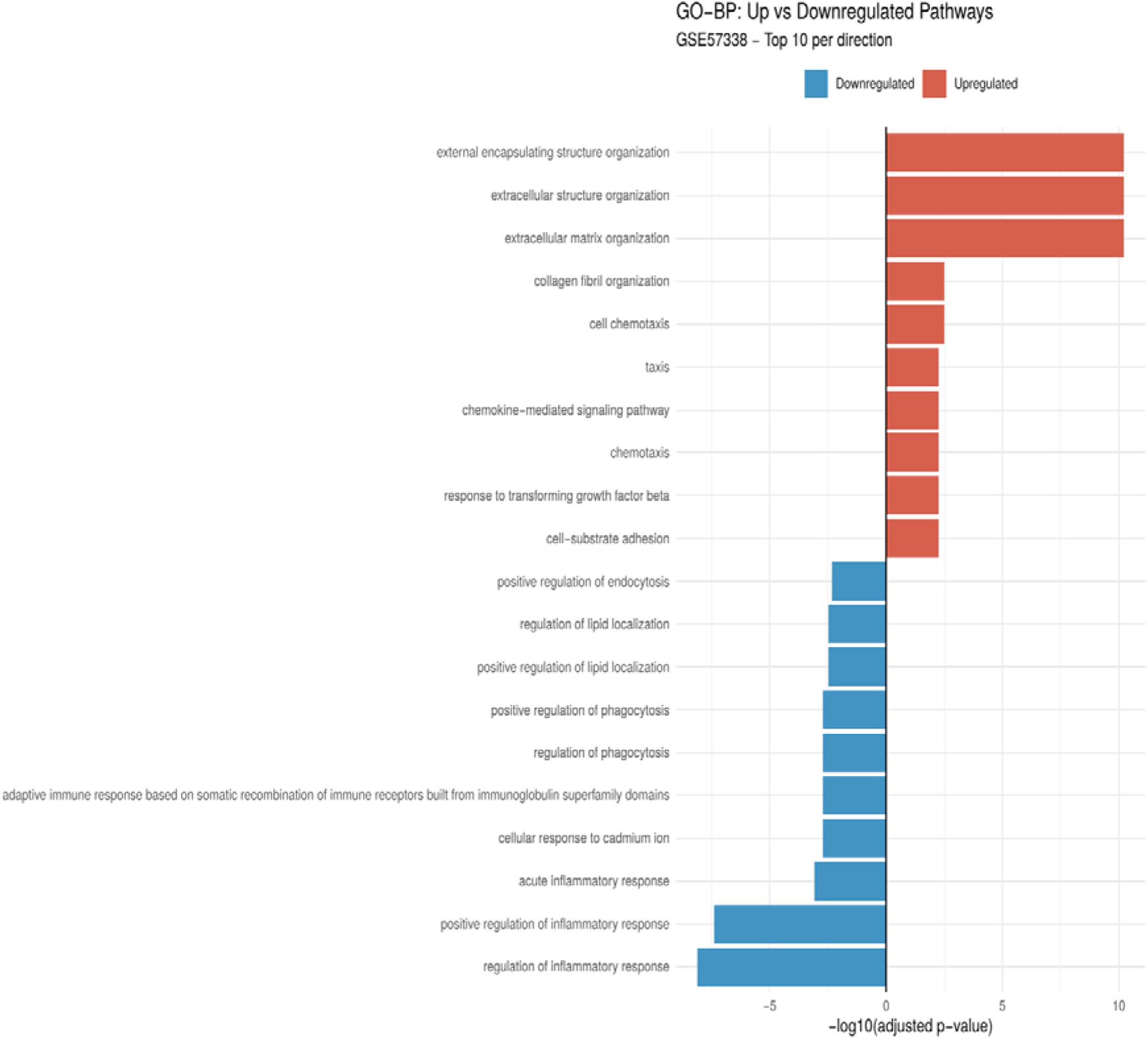
Diverging barplot comparing the top 10 enriched Gene Ontology Biological Process terms for upregulated (right) and downregulated (left) DEGs separately, showing that ECM organisation dominates upregulated pathways while immune and inflammatory response terms are enriched among downregulated genes.

**Figure 8a:**
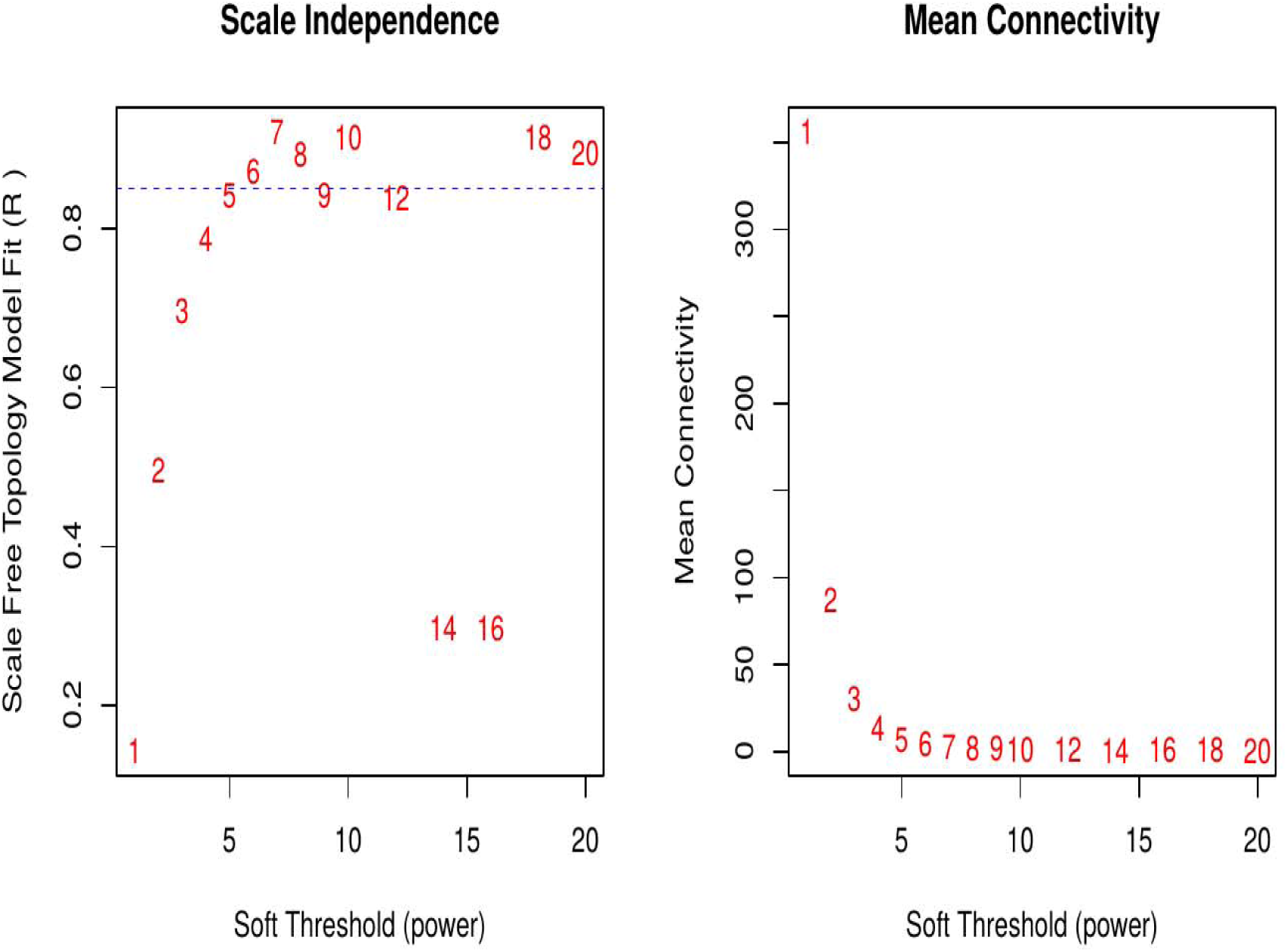
WGCNA Soft Threshold Selection. Scale-free topology model fit (R²) and mean connectivity plotted against soft-thresholding power (1-20) for WGCNA network construction in GSE57338, used to select the optimal power ensuring a scale-free network topology.

**Figure 8b:**
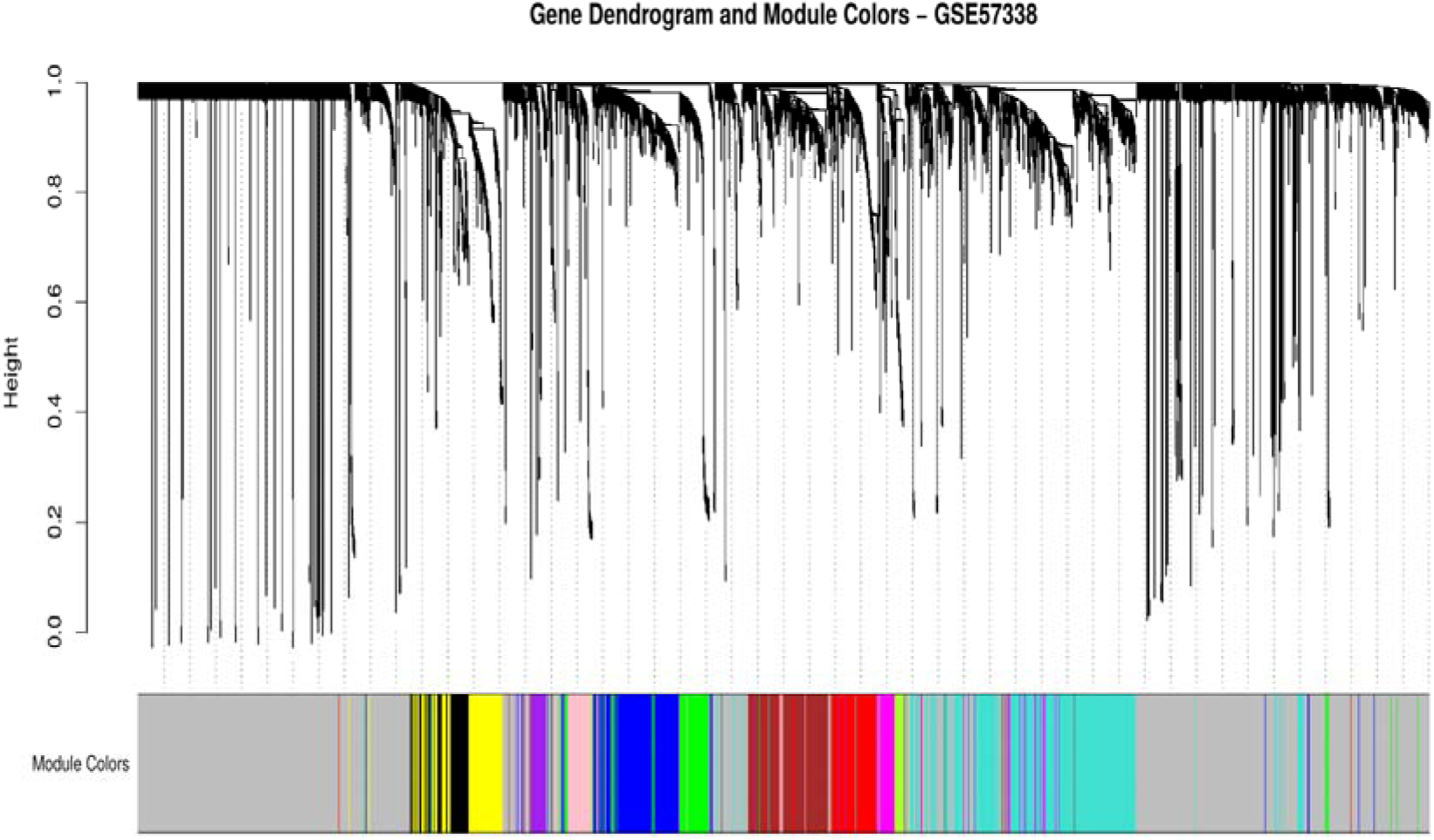
Hierarchical clustering dendrogram of the top 5,000 most variable genes (GSE57338) with assigned module colours, constructed using topological overlap matrix dissimilarity; each colour represents a distinct co-expression module.

**Figure 9:**
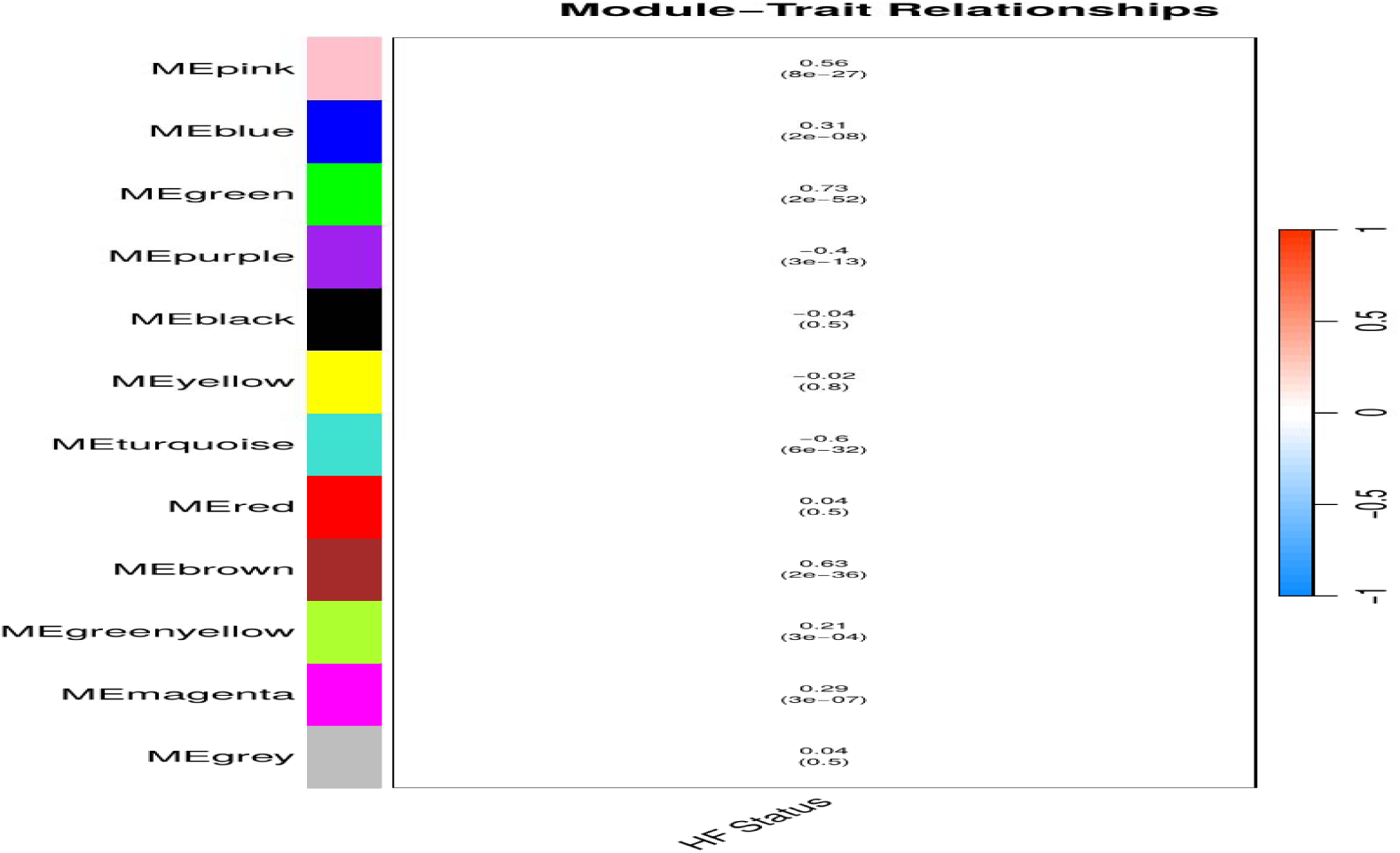
Heatmap of Pearson correlations between WGCNA module eigengenes and heart failure status (GSE57338), with correlation coefficients and p-values shown; the brown module (r = 0.63) and green module (r = 0.73) are positively correlated with heart failure, while the turquoise module (r = −0.60) is negatively correlated.

**Figure 10:**
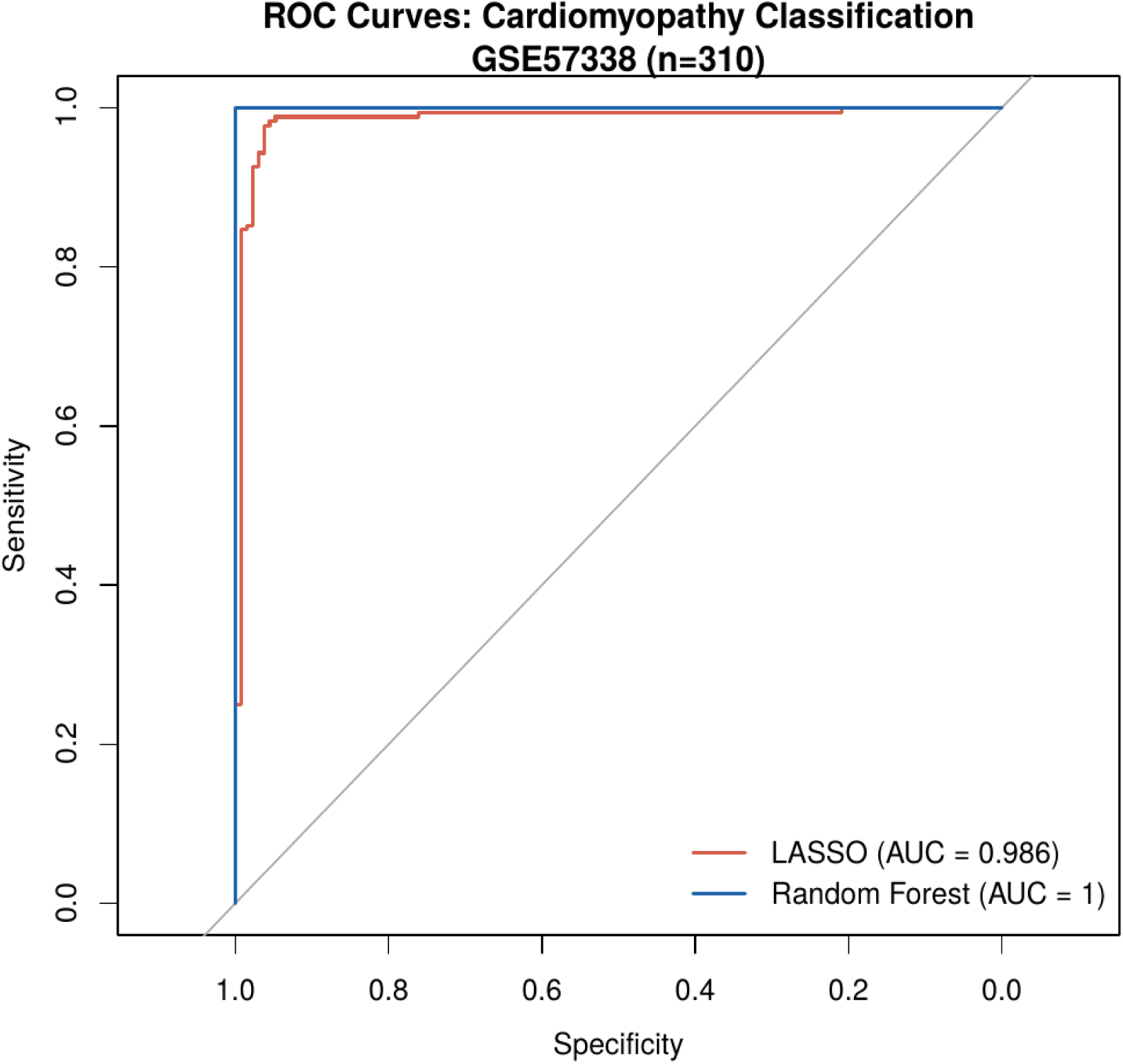
Receiver operating characteristic curves for the LASSO (AUC = 0.986) and Random Forest (AUC = 1.0) classifiers distinguishing cardiomyopathy cases from controls in the discovery cohort (GSE57338; n=310); the LASSO cross-validated AUC represents the more conservative generalisation estimate.

**Figure 11:**
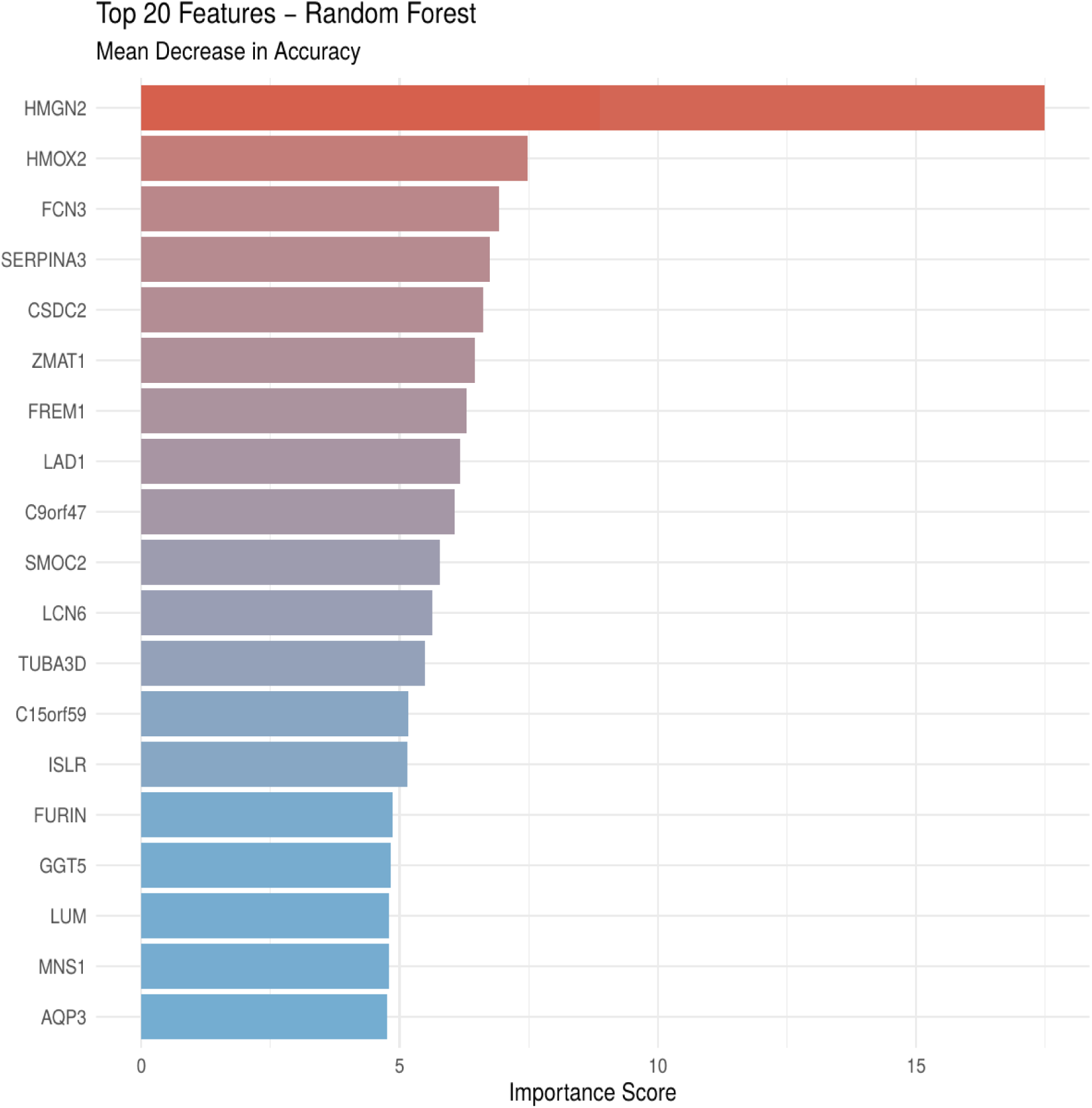
Bar chart of the top 20 genes ranked by mean decrease in accuracy from the Random Forest classifier (GSE57338), with HMGN2, HMOX2, FCN3, and SERPINA3 emerging as the most discriminating features.

**Figure 12:**
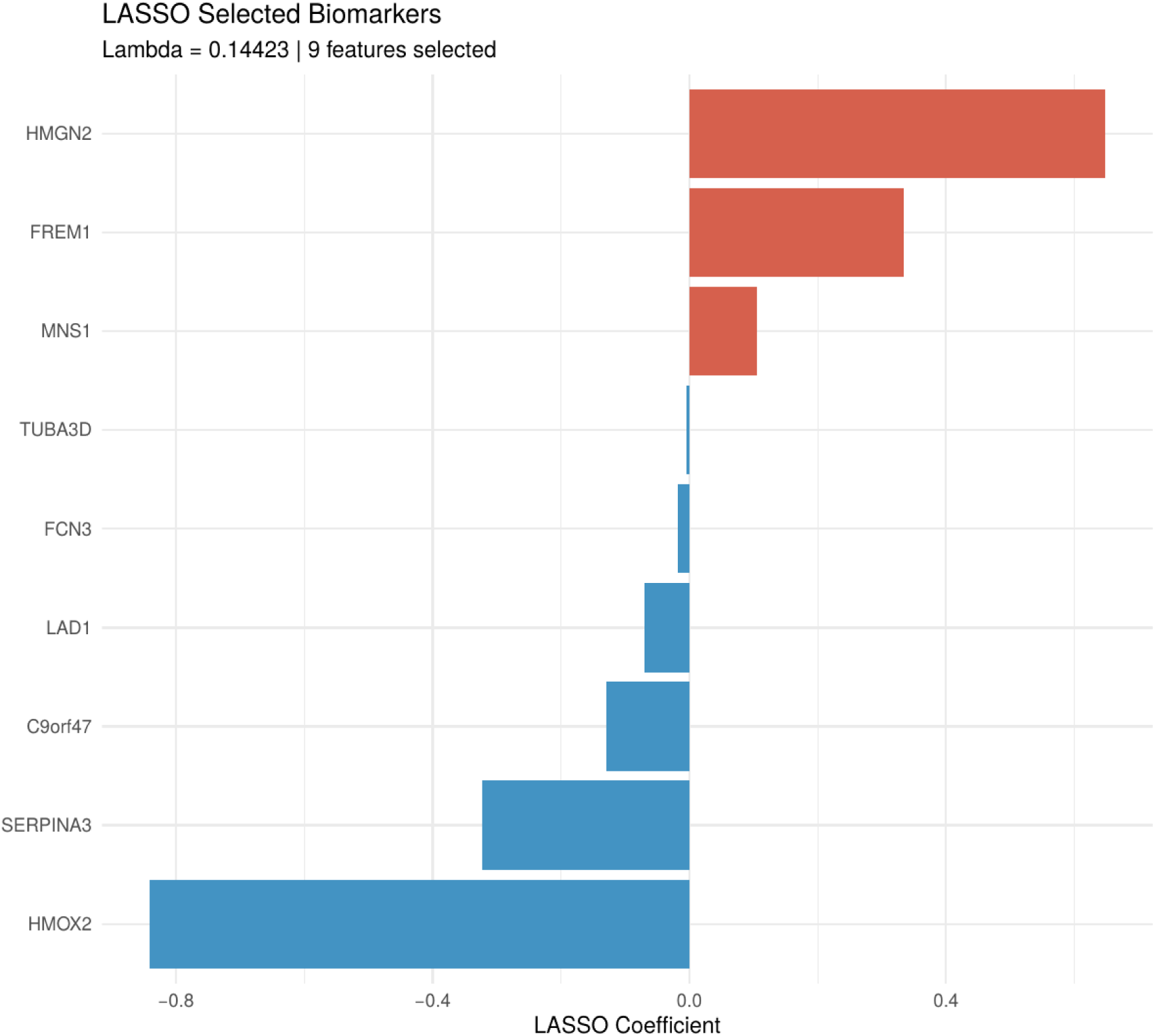
Bar chart of the nine genes selected by LASSO logistic regression (lambda = 0.144), with HMOX2 carrying the largest negative coefficient (−0.841) and HMGN2 the largest positive coefficient (+0.647), identifying them as the top discriminating features for cardiomyopathy classification.

**Figure 13:**
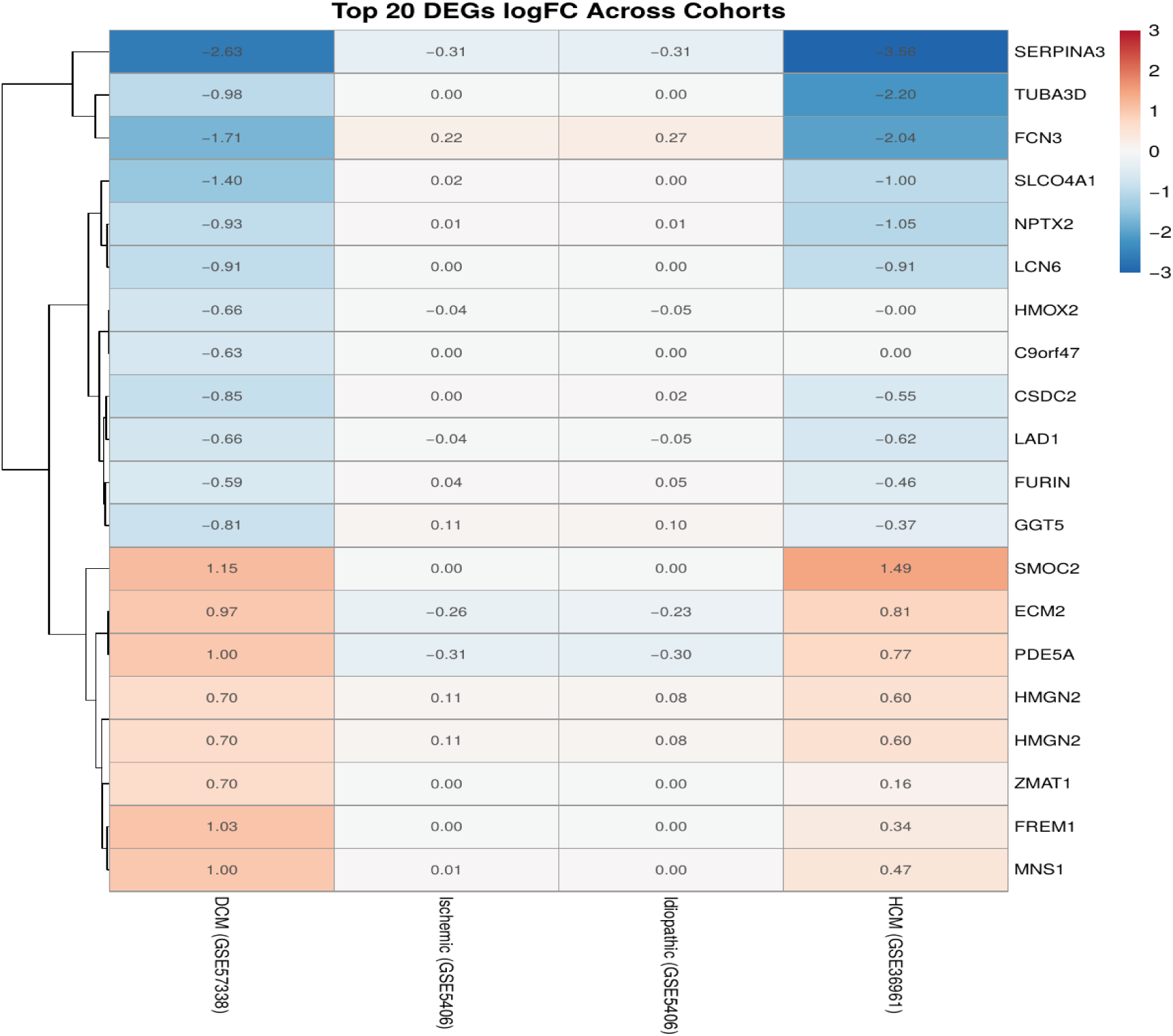
Heatmap of log2 fold change values for the top 20 discovery DEGs across four independent cohorts - DCM (GSE57338), ischemic and idiopathic DCM (GSE5406), and HCM (GSE36961) - demonstrating directional concordance for key immune and fibrotic genes including SERPINA3, FCN3, and HMOX2 across cardiomyopathy subtypes.

## Notes

### Competing Interest Statement

The authors have declared no competing interest.

